# Ultra-sensitive nanozyme-based chemiluminescence paper test for rapid diagnosis of SARS-CoV-2 infection

**DOI:** 10.1101/2020.06.05.131748

**Authors:** Dan Liu, Chenhui Ju, Chao Han, Rui Shi, Xuehui Chen, Demin Duan, Jinghua Yan, Xiyun Yan

## Abstract

The recently emerged coronavirus disease COVID-19 has now evolved into a global pandemic. Early detection is crucial for its effective control. Nucleic acid testing for viral pathogen and serological testing for host antibodies are playing important roles in current COVID-19 diagnosis. However, while nucleic acid testing is complicated, facility-restricted and time-consuming, antibody testing may result in high rates of false-negative diagnoses, especially during the early stages of viral infection. Thus, a more rapid and reliable test for both early COVID-19 diagnosis and whole-population screening is urgently needed. Here, we developed a novel nanozyme-based chemiluminescence paper assay for rapid and high-sensitive testing of SARS-CoV-2 spike antigen. Our paper test uses a newly established peroxidase-mimic Co-Fe@hemin nanozyme instead of natural HRP that catalytically amplifies the chemiluminescent signal, allowing for target concentrations to be as low as 0.1 ng/ml. Furthermore, our nanozyme-based chemiluminescence test exhibits a linear range that is 32-fold wider compared to ELISA tests. Importantly, testing is completed in less than 16 min, compared to 1-2 h required for ELISA or nucleic acid tests. Critically, signal detection is feasible using a smartphone camera. Ingredients for our test are simple and readily available, rendering overall cost considerably lower than those used in current diagnoses. In conclusion, our novel test provides a high-sensitive, point-of-care testing (POCT) approach for SARS-CoV-2 antigen detection, which should greatly increase current early screening capacities for suspected infections, and considerably lower demand for national healthcare resources.

## Introduction

Over the course of just a few months, Coronavirus Disease 2019 (COVID-19) developed into a global pandemic, causing over 370,000 deaths by beginning of June 2020^1^. Etiological investigation and gene sequencing identified a novel RNA coronavirus, provisionally named severe acute respiratory syndrome coronavirus 2 (SARS-CoV-2)^2, 3^. SARS-CoV-2 is highly infectious, with a basic reproductive number (R_0_) of approximately 3.7^4^. SARS-CoV-2 spreads through human-to-human mediated by droplets, contact or high-concentration aerosols in closed environment^5^. SARS-CoV-2 mainly infects through the upper respiratory tract and causes symptoms, such as fever, dry cough, fatigue or progressive dyspnea. One major concern for healthcare systems is discovering the early infections in incubation period and asymptomatic carriers, making a reliable test critical for successful control of the pandemic. Currently, there is no effective targeted drugs or available vaccine for COVID-19. Early diagnosis and early quarantine to cutting off the source of infection are the most effective control means of the pandemic.

Clinical features, epidemiological history, imageology and pathogenic index play a vital role in diagnosis of COVID-2019. At present, nucleic acid test based on fluorescence PCR or isothermal nucleic acid amplification are primarily employed in early diagnosis of COVID-19^6, 7^. However, nucleic acid testing requires biosafety laboratory, facility and skilled personnel, and is therefore unsuitable for field screening and out of reach for many less well-endowed communities or health care systems in developing countries. Furthermore, nucleic acid test needs RNA extraction, reverse transcription, gene amplification and result analysis, these complicated operations take at least 1∼2 h. A second major challenge associated with nucleic acid detection is the high false negative rates^8^, a potentially dangerous characteristic given the epidemiological consequences of unidentified carriers. While PCR nucleic acid testing is highly sensitive for the early diagnosis of SARS-CoV-2 infection, it is not suitable for field rapid screenings.

Antibody testing is another approach assisting nucleic acid diagnosis^9^. Colloidal gold test paper, traditional chemiluminesence immunoassay (CLIA) and enzyme-linked immunosorbent assay (ELISA) reagents have been rapidly developed^10^. Currently available antibody tests detect IgM or/and IgG antibodies against nucleocapsid or spike antigen of SARS-CoV-2 present in blood samples^11, 12^. While colloidal gold paper tests are rapid and easy-to-use^13^, the sensitivity of colloidal gold antibody test is relatively low compared with PCR-based testing. Critically, antibodies are generated only 10∼15 days past virus exposure^14^, and therefore it is not suitable for early screening and diagnosis of COVID-19, rendering early isolation and disrupting transmission impossible.

In our previous study, we reported a nanozyme colorimetric strip test for Ebola virus detection, which is based on Fe_3_O_4_ nanozyme-catalyzed chromogenic reaction^15^. Nanozymes are nanomaterials endowed with intrinsic enzyme-mimicking activity^16^, which can be used for enzyme-labeled probes. Enzymatic chemiluminescence is widely used in In Vitro Diagnosis (IVD) due to its higher sensitivity and wider linear range compared with ELISA. Traditional CLIA is based on a natural protease, such as natural horseradish peroxidase (HRP) or alkaline phosphatase (ALP), all of which are characterized by poor stability. Moreover, traditional chemiluminescence detection depends on large instruments, which are expensive and not readily available to individuals.

Here, we developed a nanozyme-based chemiluminescence paper test for rapid, high-sensitive and portable detection of SARS-CoV-2 spike antigen. We used Co-Fe@hemin nanozyme as a suitable alternative to HRP, thus combining traditional CLIA with lateral flow assay. As a novel point-of-care approach, this nanozyme-based chemiluminescence paper test will be of considerable benefit to early screening of suspected SARS-CoV-2 infections, and should greatly assist in minimizing the impact of large-scale screening on national healthcare systems, especially those in emerging and lower-income economies.

## Materials and Methods

### Materials

FeCl_3_.6H_2_O, CoCl_2_.6H_2_O, Poly (acrylic acid), Hemin, TMB, HRP, EDC, NHS, BSA, DAB and hydrogen peroxide (H_2_O_2_) were purchased from Sigma-Aldrich Co. LLC. (Shanghai, China). Polyethylene glycol was purchased from Yeasen Biotech Co., Ltd. Sodium acetate was purchased from Sinopharm Chemical Reagent Co., Ltd (Shanghai, China). Luminol substrate was purchased from Macklin Biochemical Co., Ltd (Shanghai, China). Luminunc^™^ MicroWell 96 plates were purchased from Nunc (Denmark). Nitrocellulose membrane was purchased from Sartorius (Germany). Fiberglass pads, absorbent pads and plastic boards were purchased from Shanghai Kinbio Tech. Co. Ltd. Ab1, Ab2, TP52, RP01 antibodies against SARS-CoV-2 spike antigen were provided by Academy of Military Medical Sciences and Sino Biological Inc. (Beijing, China). Pseudo-SARS-CoV-2 expressing spike S1 protein, spike S1 ELISA reagent, SARS-CoV S-RBD, MERS-CoV S-RBD, HCoV-HKU1 and HCoV-OC43 spike protein were purchased from Sino Biological Inc.

### Preparation of Co-Fe@hemin nanozyme

Firstly, Co-Fe nanoparticles (Co-Fe NPs) with carboxyl modification were synthesized through hydrothermal method. In brief, FeCl_3_.6H_2_O and CoCl_2_.6H_2_O (with a molar ratio of 3:1 to 1:1) were dissolved in 400 mL ployethylene glycol. Next, 30 g sodium acetate (NaAc) and 3 g poly (acrylic acid) were added under vigorous stirring. The solution was transferred into the high-temperature autoclave and heated at 200°C for 12∼14 h. The synthesized nanoparticles were then gathered by magnetic separation and washed with ethyl alcohol and deionized water. Subsequently, Co-Fe NPs prepared above were added into NaAc solution. Next, hemin (with a mass concentration ratio of 2.5:1) was dropwise added into the reaction system. After continuous stirring for 2 h at room temperature, the Co-Fe@hemin nanocomposites were purified by magnetic separation and washed as described above.

### Characterization of Co-Fe@hemin nanozyme

Transmission electron microscopy (TEM) images were obtained using a Tecnai Sprit microscope instrument operating at 120 KV (FEI Inc., America). The scanning electron microscopy (SEM) images were taken on a SU8020 microscope instrument (Hitachi, Japan). The hydrate particle size of Co-Fe@hemin nanozyme was measured using a 271-DPN dynamic light scattering (DLS) instrument. The UV-Vis absorbance spectra were scanned using a U-3900 absorption spectrophotometer (Hitachi, Japan) in a wavelength range of 200∼600 nm. Surface carboxyl group modification was characterized using Thermo Gravimetric-Differential Scanning Calorimeter (TG-DSC) analysis using STA449F3 Jupiter (Netzsch, German) at a heating rate of 10°C/min under N_2_ protection. The elemental analysis was performed using Energy Dispersive Spectroscopy (EDS) equipped in a SEM microscope instrument.

### Catalytic activity evaluation of peroxidase-mimic Co-Fe@hemin nanozyme

The peroxidase activity of Co-Fe@hemin nanozyme was measured according to a previously published procedure^17^. In brief, nanoparticles of different concentrations were added to 2 mL HAc/NaAc buffer (0.2 M, pH 3.6), then 100 µl TMB and 100 µl H_2_O_2_ (30% wt/vol) were added to the cuvette. The absorbance at 652 nm was measured for 1 min at the temperature of 37°C, and the rection-time curve was plotted. The specific activity (SA, U/mg) of nanozyme was calculated. The chemiluminescence catalytic activity was assessed by EnVision Multilabel plate reader (PerkinElmer) using a liquid auto-injection system and luminol substrate. The maximum chemiluminescent intensities catalyzed by Co-Fe@hemin nanozyme were measured at different pH, temperature pretreatment and different concentrations of H_2_O_2_, with HRP as positive control. Moreover, the intermediate reactive oxygen species in the reaction system were monitored by using electron spin resonance (ESR) spectroscopy and the spin trap agent, 5,5-dimethyl-1-pyrroline-N-oxide (DMPO).

### Recombinant SARS-CoV-2 spike protein and antibody preparation

The recombinant antigen is a receptor binding domain of the SARS-CoV-2 spike protein (S-RBD, located in the S1 subunit) ^18^. A cDNA sequence encoding S-RBD was expressed containing both N-terminal natural signal peptide as well as a 6×His tag at the C-terminus, after transfection into HEK293 cells. After 3 days, supernatants were collected and soluble protein was sequentially purified using HisTrap HP column and Superdex 200 column. The concentration of S-RBD protein was determined using a BCA assay kit (Pierce). The purity of recombinant protein was analyzed by SDS-PAGE. Aliquoted protein was stored at -20°C∼-80°C until further use.

Anti-Spike antibodies were generated by sorting single memory B cells as previously reported^19^. Briefly, peripheral blood mononuclear cells from convalescent COVID-19 patients were incubated with His-tagged S-RBD at 100 nM before staining with anti-CD3, anti-CD16, anti-CD235a, anti-CD19, anti-CD38, anti-CD27 and anti-His antibodies. Antigen-specific memory B cells were identified by the following markers: CD3^-^, CD16^-^, CD235a^-^, CD38^-^, CD19^+^, CD27^+^, hIgG^+^ and His^+^, and sorted into 96-well PCR plates with single cell per/well. The genes encoding Ig VH and VL chains were amplified by 5’RACE and nested PCR. The variable regions of these genes were then cloned into the human IgG1 constant region to generate full-length monoclonal antibodies, which were then expressed and purified by conventional methods. The antibody binding affinity for S-RBD was evaluated by BIAcore 8K.

### Fabrication of nanozyme-based chemiluminescence test paper

Anti-S-RBD antibody pairs were screening using ELISA method and nanozyme-based colorimetric strips^15^. The preparation of nanozyme-based chemiluminescence test paper resembles the colloidal gold strip with further modifications. The nanozyme-based chemiluminescence test paper was constituted of a PVC backing plate, an absorbing pad, a sample pad, a nitrocellulose membrane and a conjugate pad. First, the detection antibody of S-RBD (S-dAb) conjugated nanozyme chemiluminescent probes were prepared by chemical coupling of carboxyl group using EDC/NHS agents. The labeled nanozyme probes were stored in modified TB buffer containing 5% BSA. The nanozyme probes were dispensed on the pretreated conjugate pad using a BioDot dispensing equipment. The paired capture antibody of S-RBD (S-cAb) was immobilized on a nitrocellulose membrane at 1 mg/ml to form the test line (T-line), as well as the anti-human IgG antibody as the control line (C-line). Next, the above conjugate pad, nitrocellulose membrane, sample pad and absorbing pad were assembled on the PVC backing plate and followed by complete drying at 37°C for 1∼2 h. The paper boards were cut into 4 mm-wide strips. Finally, the strips were packed into clamp slots and stored at room temperature. The solid chemiluminescence substrate, excitant and dissolve buffer were stored in plastic ampoule bottles at room temperature.

### Rapid testing of recombinant SARS-CoV-2 spike antigen

In this study, rapid nanozyme-based chemiluminescence paper test of SARS-CoV-2 spike antigen was developed. Recombinant S-RBD protein of SARS-CoV-2 was used to evaluate the sensitivity of nanozyme-based chemiluminescence paper test. 100 µl of the sample containing S-RBD protein in dilution buffer was loaded onto the sample pad. The liquid sample flows through the conjugate pad and then the target analytes were recognized specifically by nanozyme probes to form immunocomplexes. Along with lateral flow, nanozyme complexes were captured by S-dAb and aggregated at T-line, presenting a brown color signal. Nanozyme probes uncombined with antigen bound with the control IgG antibody at C-line. The luminol substrate and excitant were dissolved and added onto the detection membrane after chromatography for 15min. The chemiluminescent signals of T-line and C-line were instantly captured by a smartphone camera or CCD imaging system (Clinx MiniChemi). In parallel, the sensitivity of ELISA detection for S-RBD antigen was evaluated by the double-antibody sandwich ELISA kit according to the manufacturer’s instructions.

### Rapid testing of pseudo-SARS-CoV-2

The pseudo-SARS-CoV-2 were packaged by transfecting 293T cells with a HIV-1-based retroviral vector coding SARS-CoV-2 spike S1 protein and luciferase reporter gene. The pseudovirus titer was determined as 10^10^ virus copies/ml. The spike S1 protein expressed by pseudovirus was quantified as 860 ng/ml. The serially diluted pseudovirus was detected by nanozyme-based chemiluminescence test paper according to the method described above. All virus-related operations were carried out inside a Class II biological safety cabinet.

## Results

### Characterization of peroxidase-mimic Co-Fe@hemin nanozyme

Co-Fe@hemin nanozyme is a type of nanomaterial that possesses peroxidase-mimicking activity. To characterize the morphology and microstructure of Co-Fe@hemin nanozyme, we used both TEM and SEM analysis. As shown in Fig. 1A and B, the Co-Fe@hemin nanozymes resemble spherical particles, with a mean diameter of approximately 80nm. DLS analysis showed that the mean hydrated diameter of our nanozyme spheres was approximately 100 nm (Fig. 1C). In the UV-Vis spectrum, we detected two absorption peaks for nanozyme spheres, which are corresponding to the characteristic peaks of Co-Fe nanoparticles and hemin molecules (Fig. 1D). Using the absorption spectrum, we verified that the hemin modification was present on the surface of Co-Fe@hemin nanozyme. We then analyzed the surface carboxyl group modification of Co-Fe@hemin nanozyme using TG-DSC (Fig. 1E). Furthermore, element composition and content analysis by EDS indicated that the atomic ratio of Fe to Co in the nanozyme particle is 13:1 (Fig. 1F).

**Figure 1.**
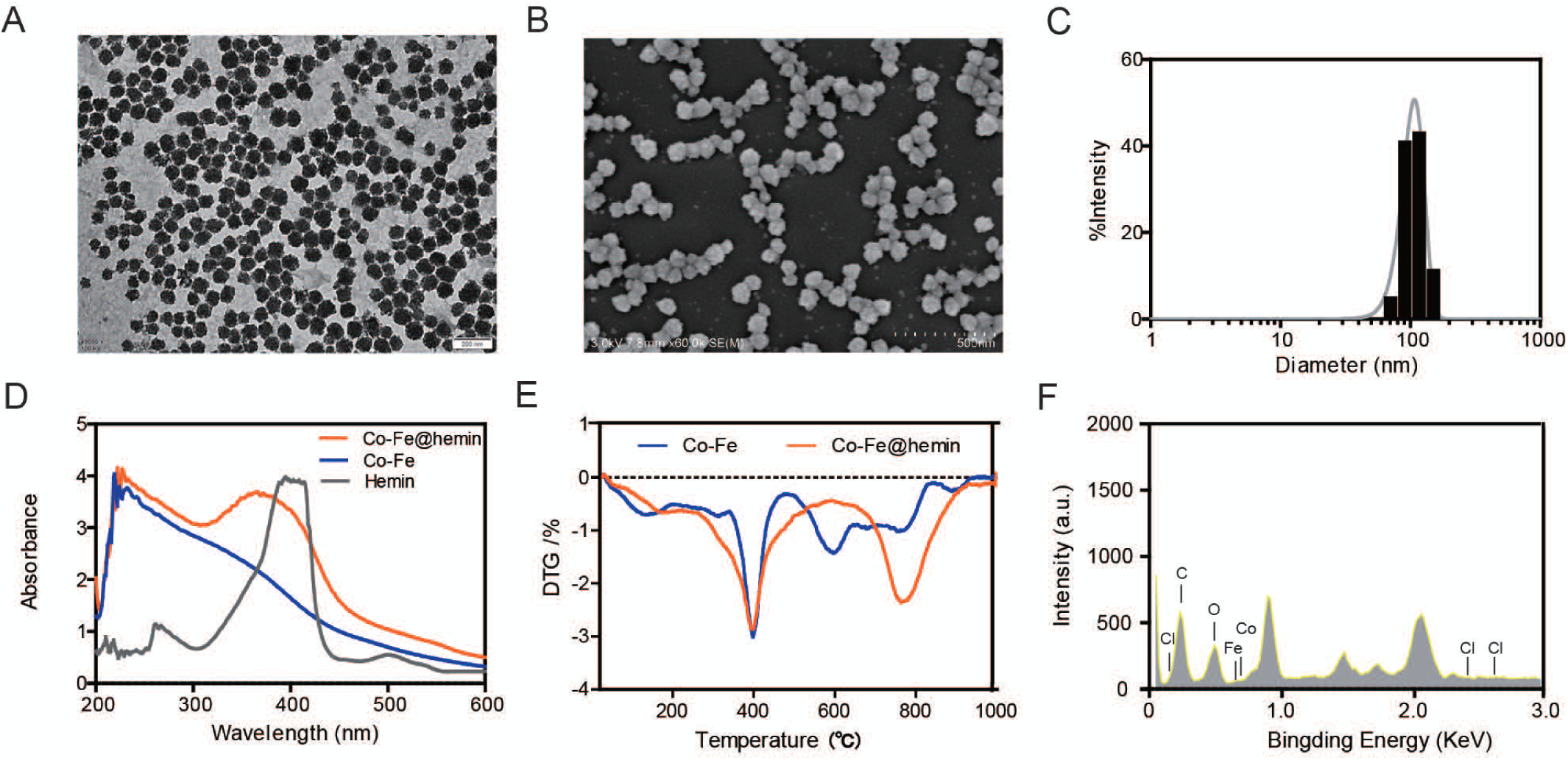
Characterization of peroxidase-mimic Co-Fe@hemin nanozyme. (A) TEM image of Co-Fe@hemin. (B) SEM image of Co-Fe@hemin. (C) Dynamic light scattering (DLS) analysis.(D) The UV-Vis absorbance spectrum of Co-Fe@hemin, Co-Fe NPs and hemin. (E) TG-DSC analysis of Co-Fe@hemin and Co-Fe NPs. (F) EDS spectrum of Co-Fe@hemin nanozyme.

### Evaluation of catalytic activity of the Co-Fe@hemin nanozyme

Peroxidase activity of our Co-Fe@hemin nanozyme was determined by the specific activity (SA, U/mg) according to the established procedures in literature^17^. TMB is one of chromogenic substrates for HRP. As shown in Fig. 2A, the catalytic activity of Co-Fe@hemin in TMB chromogenic reaction was 69.915 U/mg, which is higher than Co-Fe NPs (9.836 U/mg) and Fe_3_O_4_ NPs (5.40 U/mg, data not shown). Next, we evaluated the chemiluminescence catalytic properties of Co-Fe@hemin nanozyme for luminol substrate using a EnVision Multilabel plate reader. The chemiluminescence signals of 1 µg Co-Fe@hemin and HRP were captured in the presence of excitant H_2_O_2_. As shown in Fig. 2B, the maximum chemiluminescent intensity of Co-Fe@hemin was comparable to that of HRP, while showing a three-fold increase over than that of Co-Fe NPs. In comparison, the catalytic chemiluminescence activity of Fe_3_O_4_ NPs was found to be lower than expected.

**Figure 2.**
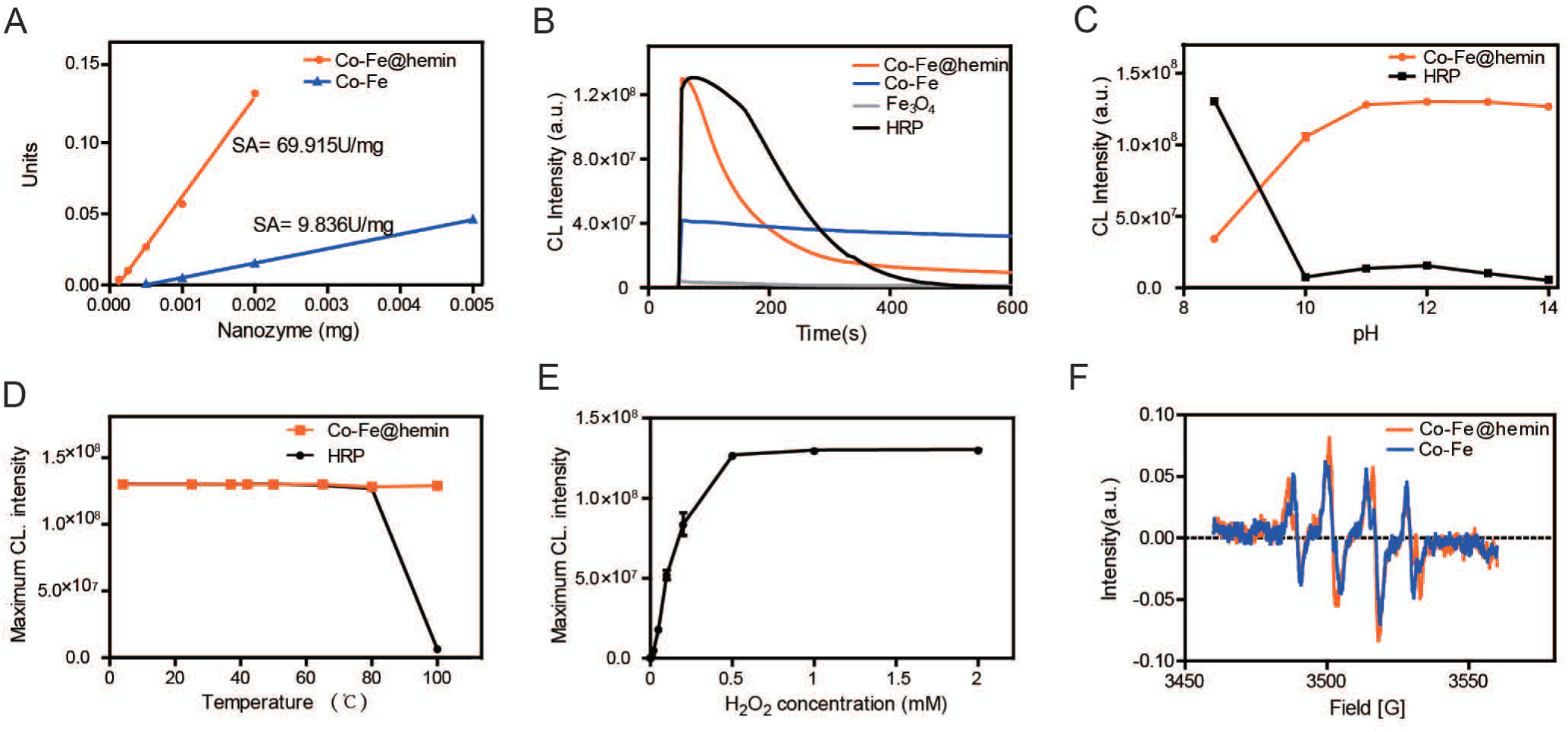
Catalytic activity of Co-Fe@hemin nanozyme. (A) Peroxidase activity of Co-Fe@hemin nanozyme in TMB chromogenic reaction. Specific activity was calculated as U/mg. (B) Chemiluminescence curve for luminol-H_2_O_2_ system. (C) Effects of pH on chemiluminescence catalytic activity of Co -Fe@hemin and HRP. (D) Effects of temperature on chemiluminescence catalytic activity of Co-Fe@hemin and HRP. (E) Effects of H_2_O_2_ concentration on Co-Fe@hemin-luminol system. Data were collected in triplicates. (F) ESR spectra of Co -Fe@hemin and Co-Fe NPs in the presence of H_2_O_2_ under alkaline conditions.

Our Co-Fe@hemin nanozyme displayed high chemiluminescent catalytic activity (the maximum intensity exceeds 8E+7) at the pH range of 9 to 14, demonstrating that our nanozyme tolerate alkaline conditions (Fig. 2C). In comparison, natural HRP only achieved its maximum catalytic activity for luminol substrate at pH 8.5, and significantly decreased as the pH increased. As shown in Fig. 2D, temperature pretreatment at 4 °C∼100 °C for 2 h failed to affect the catalytic activity of Co-Fe@hemin nanozyme. In contrast, HRP was inactivated after heat pretreatment above 80°C. Together, these results clearly showed that our nanozyme is more robust than HRP, especially under extreme PH and temperature conditions.

Next, we investigated the reaction parameters of excitant H_2_O_2_ in Co-Fe@hemin-luminol catalytic system. As shown in Fig. 2E, the maximum chemiluminescent intensity increased with the concentration of H_2_O_2_ at the range of 0.2 µM to 2 mM, reaching the plateau phase above a concentration of 1 mM. Moreover, we monitored superoxide anions (O_2_·^−^) signal in Co-Fe@hemin-H_2_O_2_ system under alkaline conditions using the ESR technique (Fig. 2F). Together, these findings indicated that luminol was oxidated by O_2_·^−^ radicals derived from the reaction of Co-Fe@hemin with H_2_O_2_, suggesting a possible mechanism of nanozyme-catalyzed chemiluminescence.

### Design of the nanozyme-based chemiluminescence paper test

After successfully validating the different core parts of our novel testing platform, we next designed the nanozyme-based chemiluminescence paper test. Fig. 3A depicts the design of our nanozyme-based chemiluminescence paper test for SARS-CoV-2 S-RBD antigen. Our paper test is based on the principle of a double antibody sandwich immunoassay. First, Co-Fe@hemin NPs labeled with S-dAb as the chemiluminescence probes were dispersed onto the conjugate pad. Along with the lateral flow of the sample, nanozyme probes combined with S-RBD antigen and S-cAb, forming the sandwich immunocomplexes. The nanozyme probes possess high peroxidase activity and thus catalyze luminol substrate in the presence of H_2_O_2_ under alkaline condition, resulting in chemiluminescence. This chemiluminescent signal is then captured and analyzed by a smartphone camera or CCD imaging system (Clinx MiniChemi). The detection can be completed within 16 min. The chemiluminescent intensity ratio of T-line to C-line is positively correlated with the concentrations of S-RBD antigen. Due to the small bonding surface of S-RBD antigen and sequence differences from other β-coronaviruses (such as HCoV-HUK1 and HCoV-OC43)^20^, it is unlikely that cross-reactivity will occur for non-target antigen.

**Figure 3.**
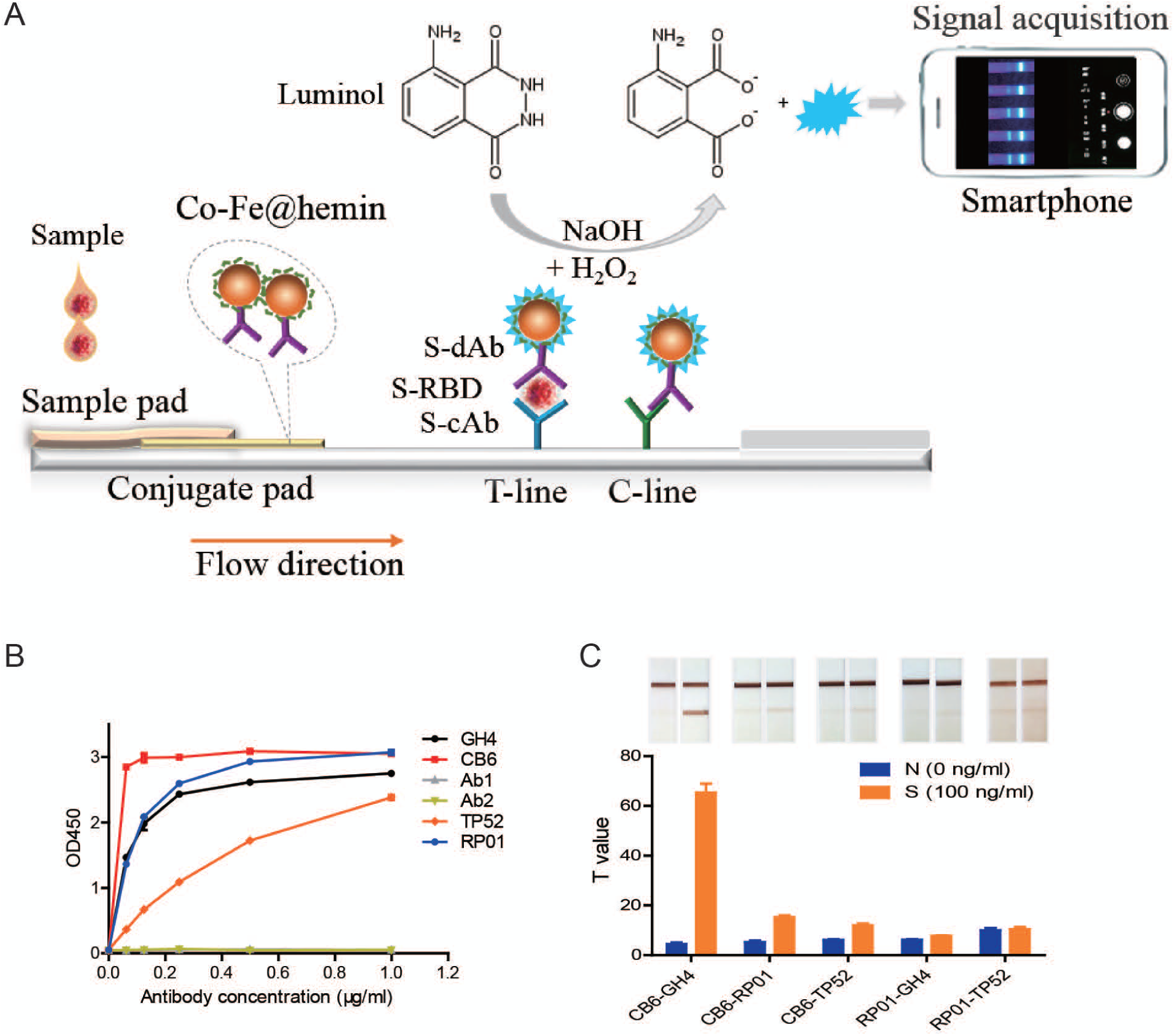
Design of the nanozyme-based chemiluminescent paper test. (A) Schematic illustration of the nanozyme-based chemiluminescence paper test for SARS-Co V-2 S-RBD antigen. Recognition, separation and catalytic amplification by nanozyme probes. (B) ELISA analysis of antibodies binding activity for S-RBD antigen. (C) Screening of paired antibodies using nanozyme based colorimetric strip. Data were collected in triplicates.

To identify the best paired antibodies against SARS-CoV-2 S-RBD antigen, we next screened a series of anti-spike antibodies using ELISA. Among the six antibodies tested, clones GH4, CB6, TP52 and RP01 exhibited high binding activity for recombinant S-RBD protein (Fig. 3B). We then manufactured the nanozyme probes labeled with CB6 and RP01 clones respectively, and combined these probes with strips immobilized with GH4, RP01, TP52 and GH4, TP52 as capture antibodies. As shown in Fig. 3C, the CB6 and GH4 pair gave the best performance in S-RBD detection. However, all other clone pairings resulted in problematic combinations, suggesting that these antibodies recognize either identical or crossed antigenic epitopes. Thus, we chose CB6 as a detection antibody and GH4 as a capture antibody for our final antibody pairs in nanozyme-based chemiluminescence paper test.

### Rapid testing of recombinant SARS-CoV-2 spike antigen

Using the CB6 and GH4 pair, we manufactured the nanozyme-based chemiluminescence test strips to detect S-RBD antigen. To assess the sensitivity of our test, we prepared a serial dilution of S-RBD protein samples. Following a chromatography procedure at room temperature for 15 minutes, the color signals of T-line were observed by naked eyes at a concentration higher than 25 ng/ml. Next, we added the luminol substrate solution. The chemiluminescent signal was instantly captured using a CCD or smartphone camera in dark environment (Fig. 4A). Next, we analyzed the chemiluminescent signal intensity using the Image-Pro Plus software. The calibration curve of antigen test was plotted as shown in Fig. 4B. Our analysis showed that our test detected as little as 0.1 ng/ml of S-RBD protein. The detection limit for S-RBD was defined as 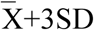 (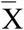: mean of the negative control, SD: standard deviation) value. The linearity of our test was identified as ranging from 0.2 to 100 ng/ml, with a correlation coefficient of 0.9855 (Fig. 4B). As the parallel test using ELISA, S-RBD antigen was detectable at 0.1 ng/ml, with a linear calibration curve ranging from 0.1 to 1.56 ng/ml (Fig. 4C). The sensitivity of our nanozyme-based chemiluminescence paper test of S-RBD protein was similar to that of ELISA, however, at least 32-fold enlargement in linearity range.

**Figure 4.**
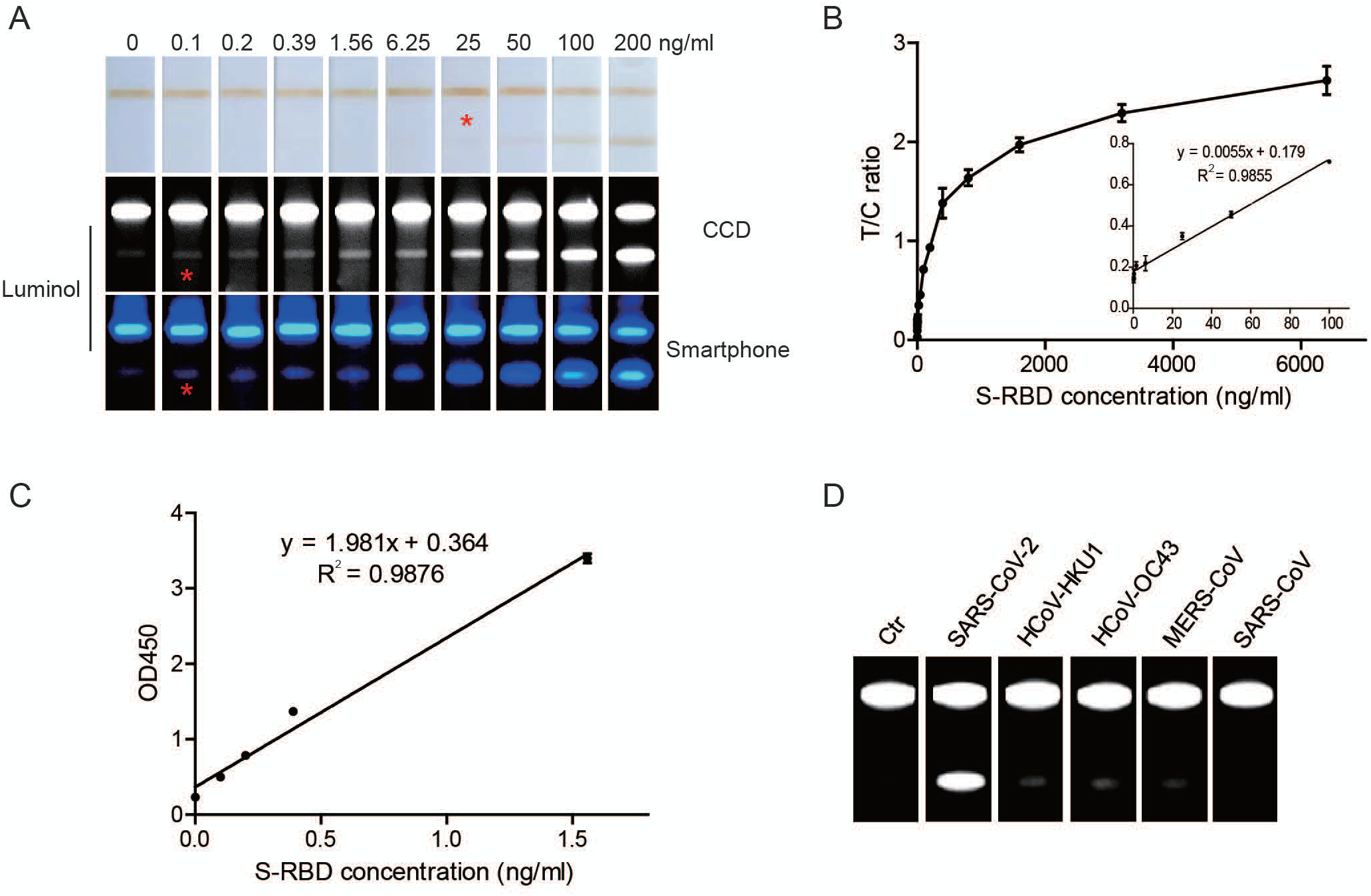
Rapid testing of recombinant SARS-CoV-2 spike antigen. (A) Gradient paper testing of SARS-Co V-2 S-RBD protein, before and after adding luminol substrate. 100 µl sample solution was loaded for each test.(B) The calibration curve of SARS-CoV-2 S-RBD detection by nanozyme-based chemiluminescence test paper. Y value was defined as the chemiluminescent intensity ratio of T-line to C-line. Concentration of S -RBD protein ranged from 0∼6400ng/ml. Samples were detected in triplicates. (C) Calibration curve of ELISA detection for S-RBD protein. Samples were detected in triplicates. (D) Our nanozyme-based chemiluminescence test paper specifically recognized the spike antigen of SARS-CoV-2, while it was negative for other human coronaviruses, such as SARS, MERS, HCoV-HKU1 and HCoV-OC43.

Moreover, the nanozyme-based chemiluminescence paper test specifically recognized the SARS-CoV-2 S-RBD protein, whereas it failed to detect the spike proteins of other known coronaviruses, such as SARS-CoV, MERS-CoV, HCoV-HKU1 and HCoV-OC43 (Fig. 4D). While SARS-CoV-2 shares more than 60% similarities with SARS-CoV, MERS-CoV, HCoV-HKU1 as well as HCoV-OC43 (cause common flu) as identified by genome sequence analysis^20^, we clearly demonstrated here that our paper test is specific for SARS-CoV-2 spike antigen.

### Rapid testing of pseudo-SARS-CoV-2

To verify the validity of our paper test in actual viral infection, we collected the pseudo-SARS-CoV-2 expressing spike protein (S1 subunit). The titer of pseudo-SARS-CoV-2 stock was determined as 1.0E+10 virus copies/ml using real-time RT-PCR. Gradient dilution series of pseudovirus were detected by the nanozyme-based chemiluminescence test paper. As shown in Fig. 5A, the detection limit of chemiluminescence paper test for pseudo-SARS-CoV-2 was 7.81E+6 copy/ml (corresponding to 0.671 ng/ml S1 protein). The linearity of our paper test was ranging from 7.81E+6 to 2E+9 copy/ml (R^2^=0.9907) (Fig. 5B). In the parallel test, the sensitivity of ELISA reached 7.81E+6 copy/ml, with a relatively narrow linear range (from 3.91E+6 to 6.25E+7 copy/ml) (Fig. 5C).

**Figure 5.**
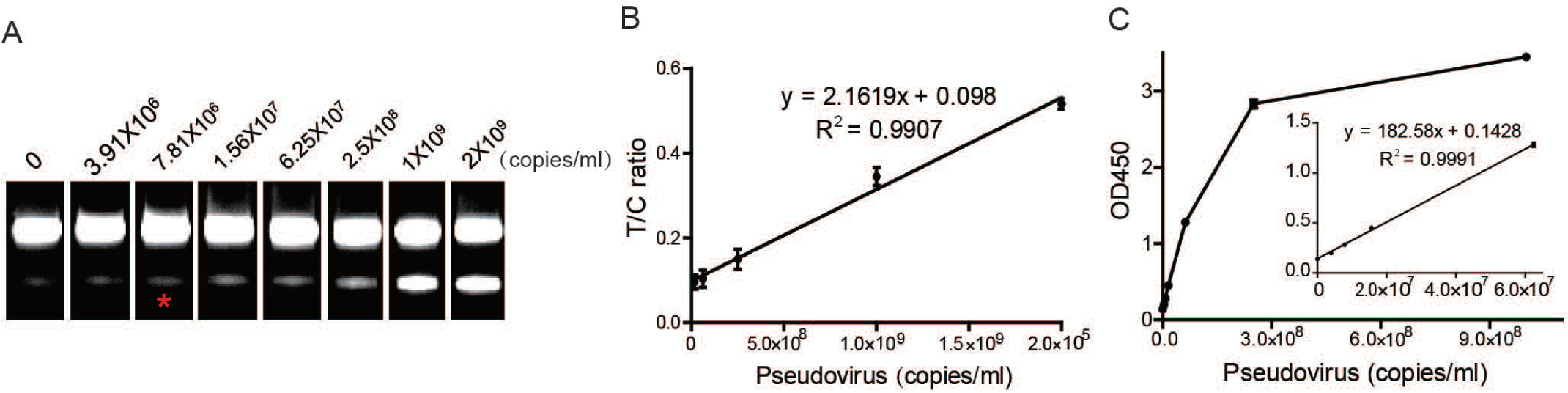
Rapid testing of pseudo-SARS-CoV-2. (A) Gradient detection of pseudo-SARS-CoV-2 samples using nanozyme-based chemiluminescence test paper. (B) Calibration curve of pseudovira paper test. (C) ELISA detection of the pseudo-SARS-CoV-2 samples. Samples were detected in triplicates.

## Discussion

Here, we present a novel SARS-CoV-2 antigen paper test that outperformed traditional paper test and HRP-based immunoassay. In this study, we successfully combined nanozyme-catalyzed chemiluminescence with a lateral immunochromatographic system to achieve ultra-sensitive as well as rapid testing for SARS-CoV-2 antigen, which should greatly benefit the early screening and diagnosis of COVID-19.

In this study, we designed and synthesized a type of peroxidase-mimic Co-Fe@hemin nanozyme that catalyzes chemiluminescence reaction as efficiently as the natural protease HRP. The sensitivity of SARS-CoV-2 antigen test by nanozyme-based chemiluminescence strip was comparable to that of ELISA. The chemiluminescence-catalyzed signal amplification makes a great improvement in sensitivity of paper test, in contrast to traditional colloidal gold, fluorescence strip or nanozyme-based colorimetric strip. The high sensitivity of detection might lower the false negative rate in the early screenings of virus infection.

In addition, our nanozyme-based chemiluminescence paper test is especially useful in POCT. Resembling traditional paper test, our test is rapid and easy to implement. Our paper test was completed within 16 minutes, which is faster than either nucleic acid testing or ELISA method. Our test can be operated without requirement for skilled personnel or dedicated site, thus it might be suitable for use at the bedside, in communities as well as open public places. Moreover, all ingredients are thermostable at ambient temperature. The detection can be performed using a portable device, thus our paper test is available and convenient for field use. In contrast, the nucleic acid and traditional chemiluminescence detection dependents on large and expensive instruments, which are only available for the central laboratory or inspection agency. Therefore, this approach for antigen test is more applicable to rapid field screening, which should greatly facilitate regional control measures of COVID-19 pandemic.

Importantly, we achieved the replacement of natural HRP using Co-Fe@hemin nanozyme in our paper test. Traditional chemiluminescence immunodiagnosis use natural proteases such as HRP or alkaline phosphatase as the core materials^21^, which are unstable, complex to produce and incur high-cost. In our study, we show that Co-Fe@hemin nanozyme exhibits better stability for temperature and acidity or alkalinity compared with HRP. Thus, the nanozyme-based chemiluminescence test paper can be stored and used at ambient temperature, benefiting outdoor settings that lack refrigeration equipment. Importantly, the catalytic efficiency for luminol substrate of Co-Fe@hemin nanozyme is comparable to HRP. In previous studies, peroxidase activities of the vast majority of nanozymes were lower than HRP. While Karyakin’s group has reported that Prussian Blue nanoparticles possess peroxidase activity superior to HRP^22^, this activity sharply decreases in alkaline condition of chemiluminescence catalysis according to our test results. The high chemiluminescent catalytic activity of Co-Fe@hemin nanozyme is key to improving the sensitivity of our paper test. Moreover, the synthesis of nanozyme is relies on readily available materials and is easily scaled up without need for expensive equipment, thus, nanozyme material is relatively low-cost compared with nature HRP, which are produced by complex extraction and purification. Therefore, the overall cost of nanozyme-based chemiluminescence paper test is relatively low compared with traditional chemiluminescence or nucleic acid testing. This advantage may reduce costs usually incurred by universal screening, and critically, benefit regions of low socio-economic standing.

While our study clearly showed that our test is superior to currently available detection procedures, it is critical to further validate the accuracy of our paper test for real clinical samples. Theoretically, this paper test will be suitable for multiple sample candidates covering throat swab, sputum, urine, saliva or serum in particular sample preservation buffer. At the time of our investigation, the officially authorized reagents for rapid testing of SARS-CoV-2 antigen remained unavailable, thus comparisons of commercial kits with our test paper in clinical sample detection require future investigation.

Lastly, our proof-of-principle study was primarily designed to enable qualitative analysis, which was achieved by means of a readily available smartphone technology. To allow for quantitative analysis of clinical samples, a quantitative test should be developed. A portable handheld device equipped with signal acquisition and data analysis function for our nanozyme-based chemiluminescence paper test is currently being developed.

In conclusion, nanozyme-based chemiluminescence paper test combines the high-sensitivity of chemiluminescence, high-specificity of immunoassay and short testing time of lateral flow chromatography technique. Due to the excellent performance, remarkable portability and low-cost, nanozyme-based chemiluminescence paper test should provide a novel point-of-care approach for early screening of SARS-CoV-2 infections. Moreover, as a universal paper-based POCT technique, it has promising potential in diagnosis application for emerging infectious diseases of pandemic potential.

## Acknowledgments

The authors would like to thank Prof. Benfen Shen from Academy of Military Medical Sciences for kind gift of anti-spike antibodies, Jiyan Zheng, Yufang Yin and Xiaonan Wang from Beijing Nanozyme Tech Co., Ltd. for technical support, Shidong Wang and Dongling Yang for great assistance. Moreover, we would like to thank T. Juelich from University of the Chinese Academy of Sciences for linguistic assistance during preparation of our manuscript. This work was supported in part by grants from the Strategic Priority Research Program of the Chinese Academy of Sciences (XDB29040101), the National Science and Technology Major Project (2018ZX10101004002004) and the Frontier Science Major Project of the Chinese Academy of Sciences (QYZDB-SSW-SMC013).

## Authors’ contributions

Xiyun Yan and Dan Liu were response for the design of the study, data analysis and manuscript preparation, Chenhui Ju, Xuehui Chen performed the detections. Demin Duan is involved in the establishment of nanozyme-based platform. Chao Han, Rui Shi and Jinghua Yan prepared the antibodies and antigen of SARS-CoV-2.

